# Serial Dependence in face-gender classification revealed in low-beta frequency EEG

**DOI:** 10.1101/2023.10.27.564174

**Authors:** Giacomo Ranieri, David C. Burr, Jason Bell, Maria Concetta Morrone

## Abstract

Perception depends not only on current sensory input but is also heavily influenced by the immediate past perceptual experience, a phenomenon known as “serial dependence”. It is particularly robust in face perception. We measured face-gender classification for a sequence of intermingled male, female and androgynous images. The classification showed strong serial dependence (androgynous images biased male when preceded by male and female when preceded by female). The strength of the bias oscillated over time in the beta range, at 14 Hz for female prior stimuli, 17 Hz for male. Using classification techniques, we were able to successfully classify the previous stimulus from current EEG activity. Classification accuracy correlated well with the strength of serial dependence in individual participants, confirming that the neural signal from the past trial biased face perception. Bandpass filtering of the signal within the beta range showed that the best information to classify gender was around 14 Hz when the previous response was “female”, and around 17 Hz when it was “male”, reinforcing the psychophysical results showing serial dependence to be carried at those frequencies. Overall, the results suggest that recent experience of face-gender is selectively represented in beta-frequency (14–20 Hz) spectral components of intrinsic neural oscillations.

**Significance Statement:** The neurophysiological mechanisms of how past perceptual experience affects current perception are poorly understood. Using classification techniques, we demonstrate that the gender of face images can be decoded from the neural activity of the EEG response to the successive face stimulus, showing that relevant neural signals are maintained over trials. Classification accuracy was higher for participants with strong serial dependence, strongly implicating these signals as the neural substrate for serial dependence. The best information to classify gender was around 14 Hz for “female” faces, and around 17 Hz for “male”, reinforcing the psychophysical results showing serial dependence to be carried at those beta-frequencies.

## Introduction

Much evidence has accumulated to suggest that the brain is a prediction machine, which generates our perceptive experience from internal models based (at least in part) on previous perceptual experience [1–4]. On this view, the brain must update predictions to minimize the discrepancy between internal models and external stimuli, in a constantly changing environment. Recent research suggest that low-frequency neural oscillations are a candidate for the role of messengers of top-down predictions [5–7].

Beta oscillations (13-30 Hz) have been linked to several perceptual, motor and cognitive processes [8–12]. Especially the lower range (up to ∼18 Hz) of beta oscillations may play a fundamental role in maintenance of working memory [13–15]. Also, they have been implicated in mechanisms of long-range communication and preservation of current brain state [16,17]. Top-down beta oscillations to macaque V1 enhance visually driven gamma oscillations [18]. Betti *et al.* [19] have proposed that beta oscillations may represent long-term perceptual *priors*. A recent study [20] reports that representations held in working memory are activated at different phases of a beta (∼25 Hz) cycle. In sum, these findings suggest that beta oscillations may be actively involved in updating priors.

Studies suggest that neural representations of recent stimuli linger in visual cortex and are boosted on the appearance of a coherent stimulus [21–24]. This signal is referred to as an activity-silent trace, as it seems not to appear in electrophysiological recordings before new stimulation. It was proposed that the underlying mechanism is a change in synaptic weights [22,24–27]. Taken together, these findings illustrate a plausible mechanistic model of prior integration, but the origins, features and dynamics of this perceptual memory trace remain unclear.

To investigate perceptual memory traces and their interaction with neural oscillations, we applied classification techniques to EEG recordings within a serial dependence paradigm. Serial dependence is a perceptual phenomenon that reveals the effects of immediate past experience on the perception of a new stimulus, integrating successive inputs on a perceptual continuum [28–31]. Liberman *et al.* [32] showed that face identities are subjected to serial dependence, whereby current perception of a face is systematically pulled towards recently seen faces [33]. Bell *et al.* [34] showed that oscillatory activity in the beta range plays an important role in discrimination of gender, and that the oscillation frequency differs between male and female images: faster for male biases (17 Hz) and slower for female biases (∼13 Hz). Based on these findings, we used a face-gender discrimination paradigm to study the neurophysiological characteristics of information of perceptual history embedded in neural oscillations. We classified previous responses across narrow-band frequency ranges, showing that the correlation of EEG signal and serial dependence peaks at separate frequencies for trials with high male and high female bias. Peak frequencies of the correlation were higher for male and lower for female images, consistent with frequencies identified in behavioral analysis.

## Results

### Behavioral results (serial dependence and oscillations in bias)

Twelve participants (5 female) were asked to classify a sequence of face images as *Male* or *Female* (Fig. 1A and methods). The images were of three types, male, female and androgynous, constructed from morphed face space. Despite calibration, aiming at 25, 50 and 75% response for female, male and androgenous stimuli, there remained a tendency to respond male rather than female to androgynous faces (average proportion male response = 0.62 ± 0.14 STD). Considering only androgynous trials, participants responded *Male* more often when the previous response was *Male* rather than *Female*, consistent with a significant serial dependence effect (Fig. 1b: average difference = 0.082 ± 0.13 STD, paired-ttest p = 0.044). The effect varied considerably across subjects, suggesting genuine individual differences, consistent with previous studies [35,36]. Separating participants based on their gender did not reveal any meaningful difference in overall bias (average proportion male response for female participants = 0.68 ± 0.21 STD, average proportion male response for male participants = 0.58 ± 0.11 STD).

**Figure 1.**
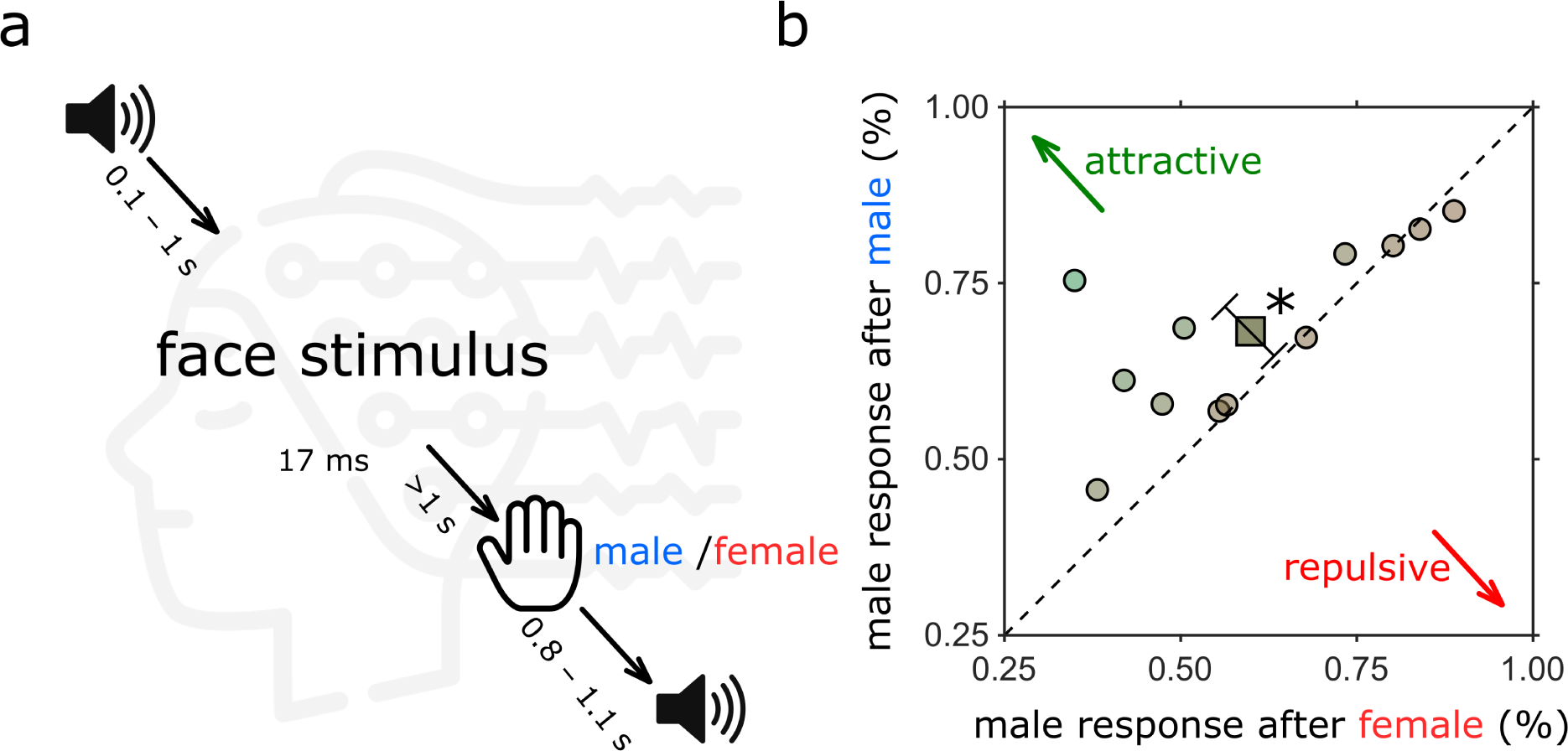
Experimental paradigm and behavioural results. **A.** Schematics of the experimental paradigm. Each trial started with an audio cue followed after a random interval (between 100-1000 ms) by a face image for 2 screen frames (16.6 ms). There were 15 different face identities, each morphed into 3 genders (androgynous, female and male), making 45 faces in total. Participants waited at least 1 s before responding by key-press, indicating whether the face appeared more female or male. A new trial started after an inter-trial interval between 800 and 1100 ms. **B.** Proportion male responses to androgynous stimuli, depending on the response to the previous stimulus (response “female” plotted on abscissa, “male” on ordinate). Yellow circles represent individual participants, with the dark yellow diamond showing the average participant. Participants above and to the left of the equality line tend to respond more male if the previous was male, and vice versa (positive serial dependence). The average participant showed serial dependence (average difference in proportion male response = 0.08 ± 0.13, p = 0.04).

As shown in Figure 1A, each trial was initiated with an auditory tone, followed by the face stimulus after an interval ranging from 100 to 1000 ms. This procedure aimed to reveal oscillations in the response, as salient auditory stimuli can reset the phase of endogenous oscillations [37]. Figures 2A & C show the oscillatory biases in responses, plotted as a function of time after the synchronizing auditory tone, separately for when the previous response was *Male* and *Female*. In both cases the responses were not constant, but oscillated over time, faster for preceding *Male* than *Female* stimuli. The red curves of Figures 2A&C show the best-fitting sinusoids, and Figures 2B&D show the associated Fourier transforms. Following a male response, bias oscillated at 18.2 Hz (p < 0.005, corrected for all frequencies in the range 10-20 Hz of the surrogate data: see Methods), and also, but less significantly at 13 Hz (p < 0.05). Following a female response, bias oscillated most strongly at 14 Hz (significance p < 0.05). These results replicate the findings of Bell et al. (2020), where bias synchronized to voluntary button press (possibly a stronger endogenous reset) oscillated at 17 Hz after for previous male faces, and 13.5 Hz for female faces.

**Figure 2.**
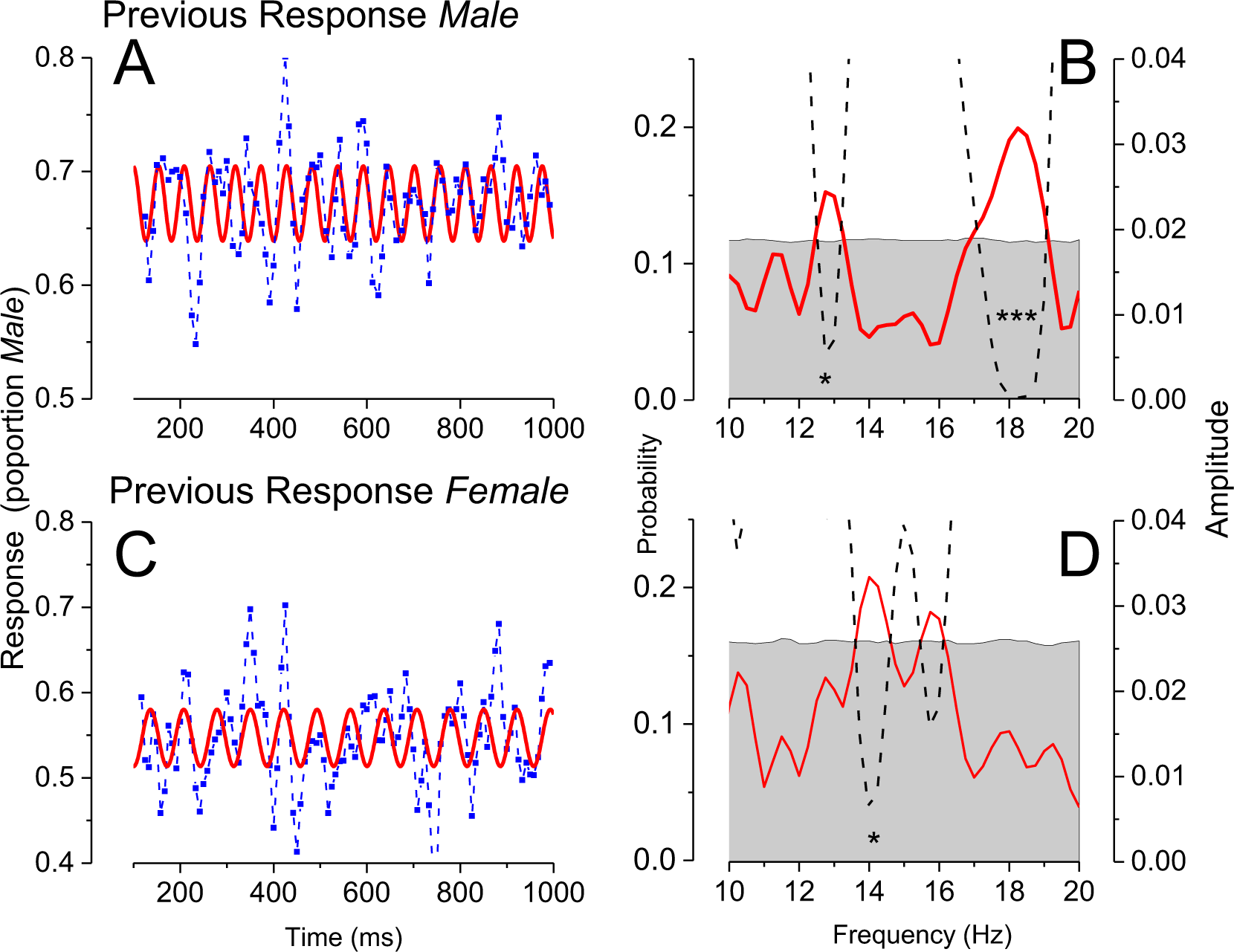
Timecourse and Fourier transforms of response bias. **A.** Proportion of *“Male”* responses to androgynous stimuli preceded by response *Male*, as a function of time after the auditory tone (blue symbols). The red curve shows the best fitting sinusoid, at 18.2 Hz. **B.** Fourier transform of the data of Fig. A, showing the strongest peak at 18.2 Hz. There is a secondary, weaker peak at 13 Hz. The dashed black lines show probability of significance, corrected for all frequencies in the range 10-20 Hz of the surrogate data **C.** Same as A, but for stimuli preceded by response *Female*. Best fitting sinusoid at 14 Hz. **D.** Same as B, for stimuli preceded by response *Female*.

### EEG results (ERP and power analysis)

The main purpose of this study was to study the neural mechanisms behind serial dependence, recording EEG from participants while they made sequential psychophysical judgements. We first analyzed the ERPs for the two conditions “previous response male” and “previous response female” (average result in Fig. 3 a-c for electrodes Fz, CP5 and Oz). There were no clear or significant differences between the two conditions. We also analyzed the responses aligning them to the phase-resetting auditory tone (Fig. 2 d-f), but again, there were no significant differences in the responses.

**Figure 3.**
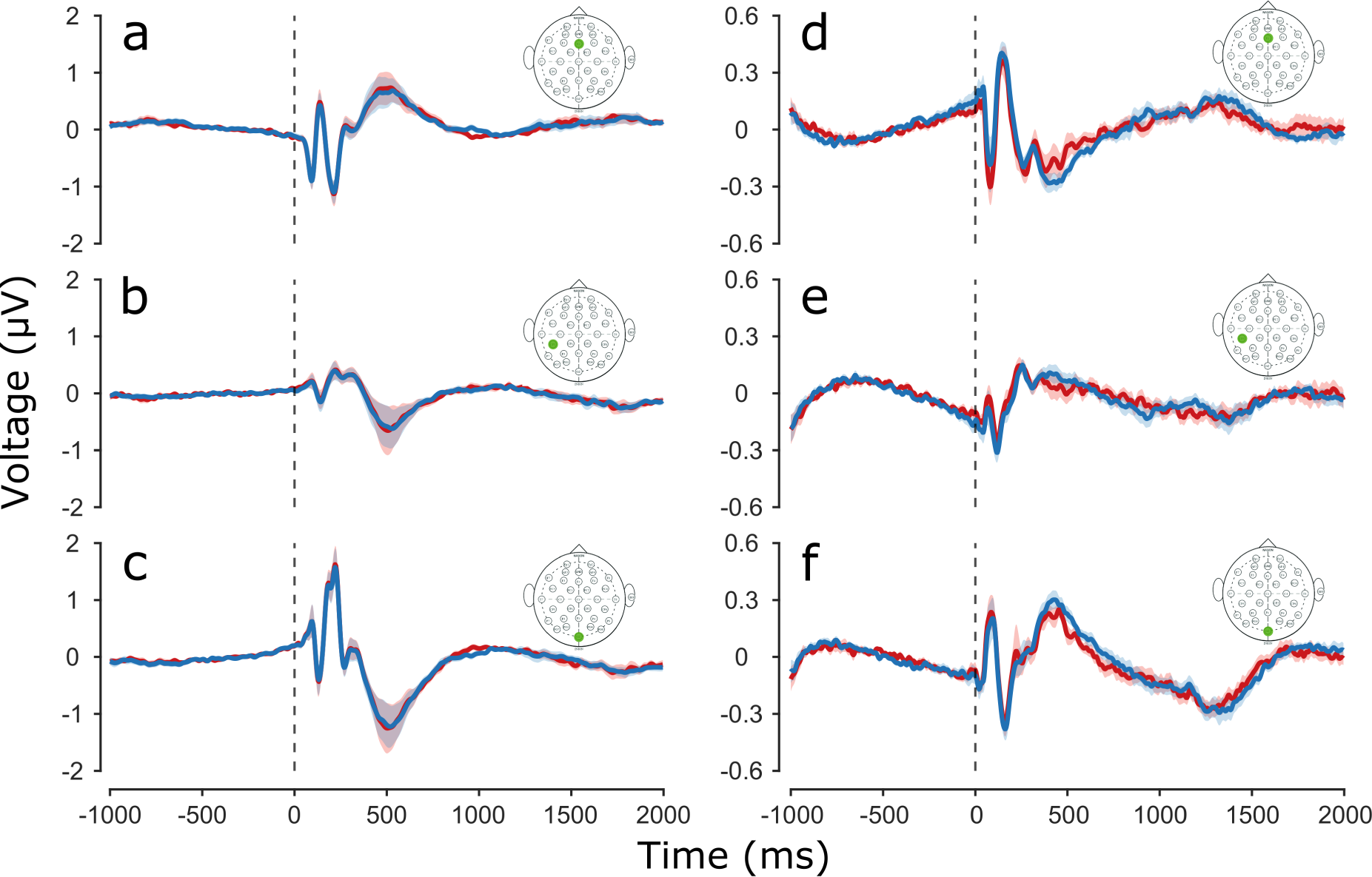
Event-related potentials (ERPs) separated according to previous response. **A-C.** ERP synchronized to face stimulus presentation of example Fz, CP5 and Oz electrodes. In blue ERP of trials where participants responded male to the previous trial, in red where they responded female. Color shading represents ±Standard Error of the Mean (SEM). **D-F.** ERP synchronized to audio cue presentation of Fz, CP5 and Oz electrodes.

### Decoding results

We next used classification techniques to test whether decoding EEG signals to the current trial could classify the responses to the *previous* face stimuli. This technique is based on small differences in signal distribution across electrode position. It relies on few assumptions, as all brain activity is used, without selecting electrodes or ROIs.

Figure 4a shows the main result. It shows accuracy of decoding the *previous* psychophysical response from the EEG response to the current stimulus, as a function of time after (current) stimulus onset. The curve is consistently above chance (0.5, for the two possible responses), and reaches the stringent significance level between 340 and 560 ms after stimulus onset, and also between 1080 and 1200 ms. This shows that there is information about the *previous* trial in the neural response to the current trial. Figure 4b is the temporal generalization matrix showing classification accuracy across all training and testing times. The decoding performance of the trained models for the specific time interval is plotted as a heat map. There are three regions where decoding was significantly above chance (within the white regions of Fig. 3b), two corresponding to those illustrated in Figure 4a (which is the diagonal of the matrix), and an extra one at 0 training and about 120 ms testing. The regions of high accuracy are distributed mainly across the diagonal, but the additional significant region away from the diagonal is evidence for some generalization for different training and testing times.

**Figure 4.**
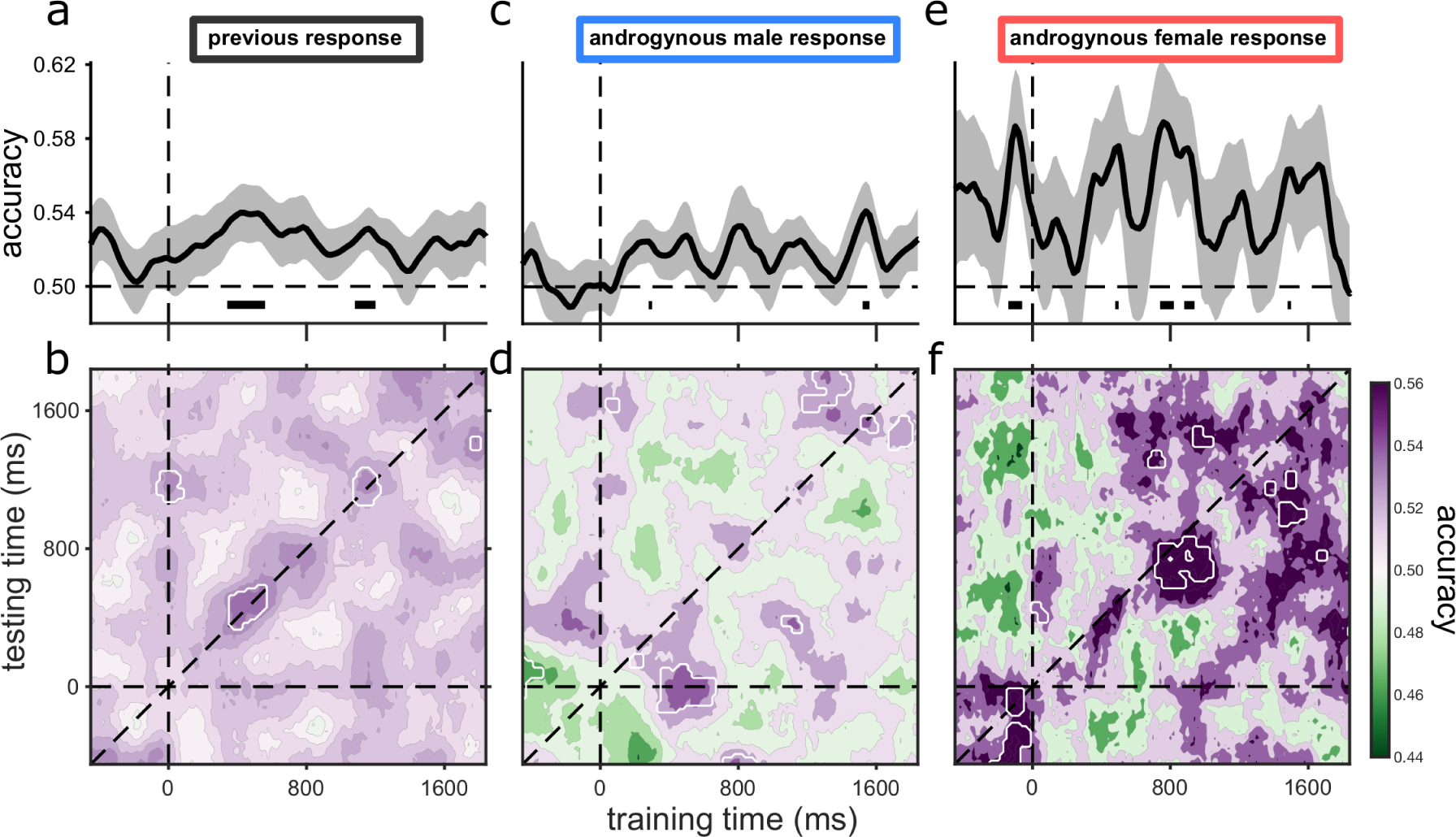
Decoding of previous responses from current-trial EEG activity. In all decoding procedures, classifiers were trained to distinguish between previous response male and previous response female, on all trials. **A.** Classifier accuracy as a function of duration after trial onset for all trials of the previous stimuli. **B.** Generalization matrix for the same stimuli, decoding all possible combinations of training and testing times. White contours indicate regions of significant accuracy after cluster correction. Decoding was strongest along the diagonal (which corresponds to figure A), but there were also significant regions of decoding away from the diagonal, suggesting generalization of decoding. **C,D** Same as A & B, but only when the current response was male and the previous stimulus androgenous. **E,F** Same as A & B, but only when the current response was female and the previous stimulus androgenous.

To investigate further the relevance of this trace on participant responses, we considered only trials where the current stimulus was androgynous, and further separated them on the basis of participant response to it (male or female: Fig. 4c–f). We took classifiers trained on all *previous* trials (as for Fig. 4a&b), but tested them on these two subsets of data: current androgenous stimuli with response *Male* and with response *Female*, to examine separately decoded signal in trials where participants had higher likelihood of being biased towards one of the two responses based on the presence of the memory trace.

The results show that previous response traces are much more readable in androgynous trials where participants responded *Female* (*difference in peak accuracy* = 0.12 ± 0.03 SEM, *p* = 0.003, Log(BF10) = 1.16). This is possibly due to the overall tendency of participants to respond male, so there is more information in a response *Female*. Again the highest accuracy tends to be on the diagonals, but some regions off-diagonal are significant, pointing to limited generalization of coding and decoding.

Activation maps (Fig. 5a) show the time-course of which electrodes were most informative for classification. Before stimulus presentation decoding relied on a stable right-occipital dipole and on distributed frontocentral locations. In early perceptual processing (up to ∼60 ms) the memory signal was classified by activity of occipital electrodes. The signal shifted progressively to frontal locations up to 140 ms, when just frontal electrodes contributed to classification. At 300 ms, occipital electrodes started to contribute again to the signal. At around 700 ms parietal and frontocentral locations became relevant, remaining relatively stable up to about 1600 ms. Activation maps before stimulus presentation and well after stimulus presentation were quite different, consistent with the lack of generalization for those intervals across training and testing time (Fig. 4 b,d,f top-left and bottom-right corner).

**Figure 5.**
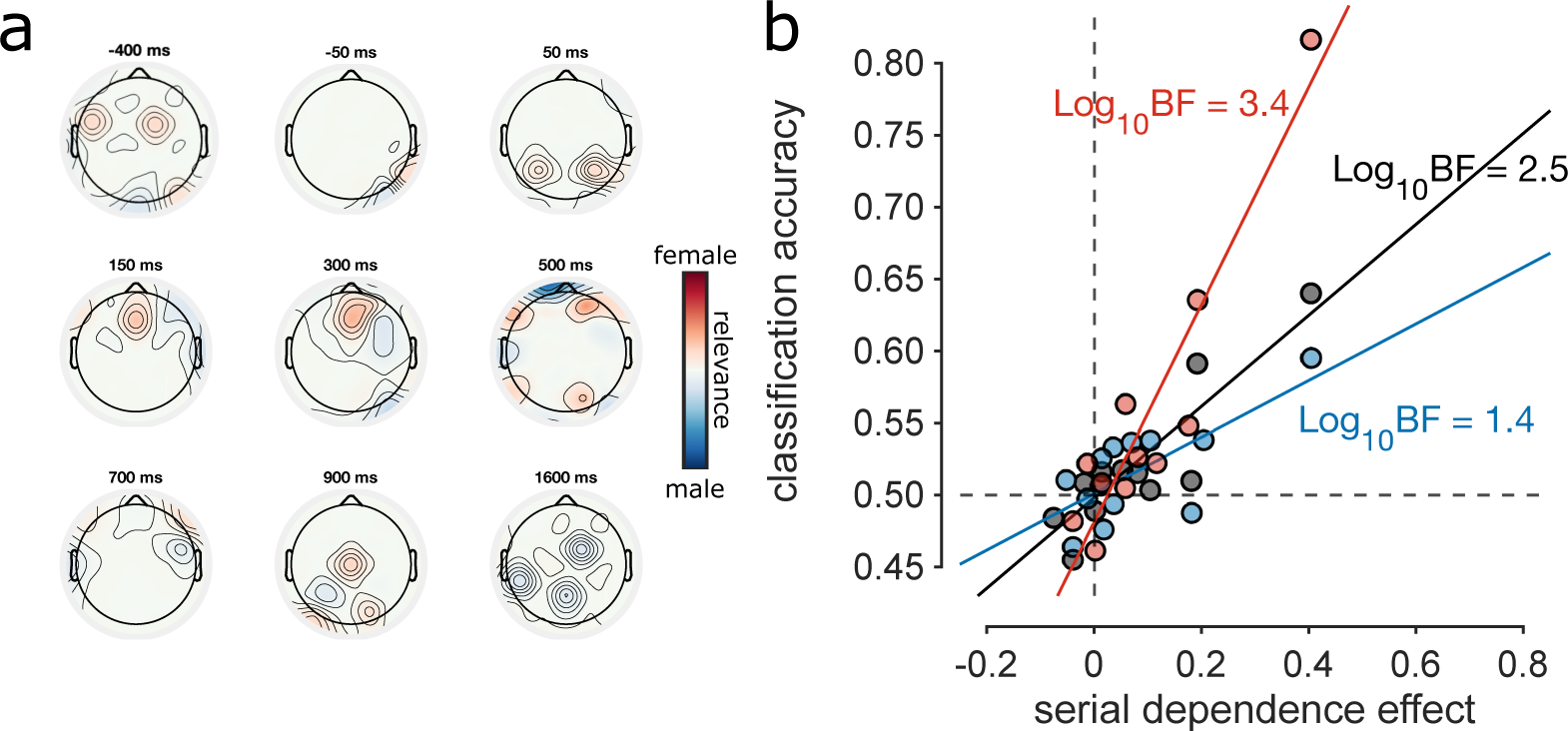
Activation patterns and correlation with behavior. **A.** Activation patterns of classification showing the relative weights assigned to electrodes for decoding for example time points. Positive weights (red) indicate that higher power at the location sways classifiers to identify the signal as belonging to the “previous female” class, while negative weights (blue) indicate that higher power sways classification towards “previous male”. Weights are normalized on maximum activation across electrodes and time. **B.** Correlation between serial dependence strength and classification accuracy across participants, for the 3 decoding conditions in Fig. 3. Serial dependence strength (abscissa, constant across all 3 conditions) was calculated as the difference between proportion male response with previous male response and proportion male response with previous female response. Classification accuracy was calculated by averaging individual participant accuracy across the diagonal ± 2 points (final precision ±60 ms) in the window 0–1 s. Decoding tested on all trials in black, on androgynous trials with female response in red, on androgynous trials with male response in blue. The strength of the correlation is given by Log_10_ Bayes Factor, shown near each fit.

We tested the correlation between classification accuracy and serial dependence across participants (Fig. 5b). Classification accuracy was averaged over the entire post-stimulus time window of the temporal generalization matrix, and serial dependence calculated as the difference of proportion male responses on androgynous trials when the previous response was male compared with when it was female. We found a significant positive correlation of serial dependence with average decoding of all trials (black dots, *r* = 0.89, *p* < 0.001, Log10BF = 2.5). The correlation of memory trace and serial dependence was stronger for classification accuracy of female-biased trials (red dots, *r* = 0.93, *p* < 0.0001, Log10BF = 3.4) compared with male-biased trials (blue dots, *r* = 0.80, *p* = 0.002, Log10BF = 1.4, but the difference was not statistically different (*rdifference* = 0.13, *p* = 0.23).

To relate the EEG results to the psychophysics, which showed clear beta-frequency oscillations, we repeated the decoding analysis after filtering the EEG into narrow-band windows, from 4 to 20 Hz. Figure 6A shows how decoding accuracy varied with filter frequency. Considering all androgenous stimuli, average classification was relatively flat across frequency (black dashed line, Fig. 6a). However, confining the analysis to androgynous stimuli with female response shows slightly decoding around low-beta, 12-17 Hz (red line, Fig. 6). The trend with androgynous stimuli with male response was less clear, but tended to increase over the beta range, peaking at the maximum analyzed, around 20 Hz. This is consistent with the psychophysical results, but not strong support.

As decoding accuracy covaried with the magnitude of serial dependence, showing a strong link between the psychophysics and EEG, we next tested whether the strength of the correlation my vary with frequency range. Figure 6b shows how the square of the correlation coefficient (R2, the variance explained by the correlation) varies with filter frequency. The correlation considering all androgenous trials are again relatively flat, but the correlation for the androgenous stimuli with *Female* response shows peaks at around 14-15 Hz, while that for androgenous stimuli with male response peaks at 17 Hz. Again this is very similar to the psychophysical results showing peaks at 14 and 18 Hz for previous *Female* and *Male* responses. Figure 6C shows the log Bayes Factor associated with the correlations, showing even clearer peaks at 14 Hz for *Female* and 18 Hz for *Male*.

## Discussion

This study investigated whether neural endogenous oscillations may be instrumental in transmitting predictive information about face-gender. The results show that previous responses leave a lingering EEG trace that can be decoded during the processing of a new stimulus. We report strong correlations between the strength of behavioral serial dependence effects and the strength of neural classification of previous responses. Importantly, the correlations peak at similar frequencies to those identified in behavioral analysis, both in previous research [34] and here, suggesting that perceptual representations of recent experience may be encoded within the spectral structure of neural oscillations.

Since classifiers are trained on averages of single-trial EEG amplitude envelopes, decoding could rely on both phase-locked and non-phase-locked information. Temporal generalization maps reveal sparse regions of significant decoding of the previous response, but exploring the distributions of accuracy, especially for androgynous trials with female response, suggests that regions of high accuracy may be distributed over the entire temporal matrix, rather than being confined to the diagonals where training and testing time coincide. This qualitative interpretation would be consistent with a memory signal that is relatively stable in time, with discrete evolution at specific time points [38]. Decoding accuracy was strongest for androgynous trials with female response, and weakest for androgynous trials with male response. This may be explained by the overall male bias of participants, so the female response was more informative.

To demonstrate decoding of previous trials, we chose to use responses rather than stimuli as class labels, for several reasons. Firstly, on half the trials the stimuli were androgynous, neither male nor female, and therefore uninformative: yet the response to those stimuli is very informative, reflecting an internal state rather than the stimulus. Secondly, as responses to male and female stimuli were about 75% correct, stimuli and responses are highly correlated, hard to disentangle. We are aware of the ongoing debate on whether serial dependence acts on early perception or on later decision-making processes [39–43]. However, responses do not represent only late decision stages, but the internal neural state of the participant, so the choice of using responses does not speak to this issue. Inspection of the activation patterns of decoders shows that classification relies mainly on occipital electrodes during early stages of visual processing. This is consistent with quick activation of early visual cortices reported in previous studies on serial dependence [41,44–46]. The immediate activation of visual areas, as early as 50 ms after face presentation (earlier than we would expect for bottom-up visual processing) suggests that the representation of the prior may be already embedded in visual cortices, possibly in the form of synaptic gain changes [22,23,47]. Lastly, the correlation between neural signal and behavior and the coherence in frequency found in the GLM analysis points to beta oscillations as the generator of the behavioral bias.

Recent studies have employed classification techniques to reveal traces of previous trials, suggesting that perceptual experience lingers as an activity-silent trace, which is then reactivated with new stimulation [21–24]. In these paradigms, classification accuracy is at chance level before the presentation of a new stimulus, and significant decoding is reported only after. However, researchers have also questioned the activity-silent trace hypothesis. Runyan *et al.* [48] identified replay of neural activity during rest, which may be related to ongoing consolidation processes that help to strengthen memories over time. Stokes *et al.* [49] also challenge the idea, suggesting that perceptual learning is instead supported by changes in neural connectivity and plasticity. We have previously supported the activity-silent trace hypothesis [24], finding that classification accuracy (for spatial frequency discrimination) was not significant before new stimuli were presented. However, the temporal generalization matrices (Fig. 3) for the female bias condition show that the trace is sometimes present before the presentation of a new stimulus However, in interpreting these results, we also have to consider that the generalization matrices for female bias is noisier, given the lower number of trials available. Furthermore, classifiers trained before time 0 do not accurately classify previous traces after time 0, and vice versa. This suggests that there is a distinct change in the signal when a new stimulus is presented, supporting the action of a silent memory signal.

It is widely acknowledged that observers enhance efficiency by using past information to anticipate future sensory input. The connection between behavior and decoding of previous responses was notably pronounced when limiting the neurophysiological signal to low-beta frequencies, where the behavioral bias oscillated according to the previously perceived gender. It has been established that low-frequency oscillations contribute to the transmission of predictive information, akin to the concept of perceptual echoes put forth by VanRullen & Macdonald [50]. Ho *et al.* [51] showed that auditory stimuli oscillated within the alpha range (∼9 Hz) at distinct phases when presented to either the left or the right ear. Using a similar paradigm to the present study, Bell *et al.* [34] demonstrated that following the observation of a specific stimulus, whether male or female, the inclination to perceive an androgynous face as female or male oscillated respectively at 13.5 or 17 Hz. Considerable evidence exists detailing how bias exhibits oscillatory behavior at specific frequencies, as observed in visual orientation discrimination [36], trans-saccadic location discrimination [52], audiovisual temporal judgement [53]. For more intricate perceptual functions, like face gender discrimination, it is possible that prior expectations require a coding mechanism involving multiple frequency channels. The literature suggests that beta oscillations may play a role in processing local features [54,55]. It is conceivable that various frequencies of beta oscillations might explain the response patterns elicited by female or male stimuli, as well as more complex stimuli in general. The arrangement of local facial features could potentially clue the interpretation of faces as masculine or feminine.

Overall, our results suggest that recent experience of face-gender is represented in low-frequency spectral components of intrinsic neural oscillations (low-beta 14–20 Hz). The strength of the active trace correlates with the strength of serial effects in behavior, especially in the low-beta range, suggesting that our signal may be the underlying neural substrate of the attractive effect. These results suggest that recent experience lingers in perceptual cortices and changes with new stimulation, possibly becoming strengthened. The strong correlation between decoding accuracy and the strength of serial dependence further suggests that these oscillatory signals are highly instrumental in the transmission of internal models, within the predictive coding framework [5,56]. Overall, our results corroborate the intuition of Bastos *et al.* [57] that in a hierarchical predictive coding framework, low-frequency neural oscillations in encephalography (4–22 Hz range) are a good candidate for top-down internal models.

## Materials and Methods

### Participants

Twelve healthy adults (7 females, age range 20 – 29 years, mean = 24.8 years, SD = 2.7 years) participated in the experiment with monetary compensation (10 €/h). All participants had normal or corrected-to-normal vision and gave written informed consent. The experimental design was approved by the local regional ethics committee (*Comitato Etico Pediatrico Regionale — Azienda Ospedaliero-Universitaria Meyer — Firenze*), and is in line with the declaration of Helsinki for ethical principles for medical research involving human subjects (DoH-Oct2008). We did not perform a formal power analysis to determine participant number but, based on our previous experience with decoding EEG [24] and also psychophysically measured oscillations of face gender [34], we reasoned that 12 should be sufficient.

### Stimuli and apparatus

The experiment was recorded in a quiet dark room, where participants sat in a comfortable chair with head rested on a chin rest. The stimuli were presented on a Display++ LCD Monitor (Cambridge Research Systems, 120 Hz, 1920 x 1080 resolution), gamma corrected, 70 cm from the eyes, mean grey screen luminance equal to 50 cd/m^2^. Face stimuli were a subset of images taken from Bell et al. (2020). They were originally generated in FaceGen Modeller 3.5.3 and saved as high resolution 2D grey scale image (6.6° x 6.6°). The faces were white, mid 20s, with gender neutral coloring, shape, and typical asymmetry. We performed a preliminary response-balancing procedure on 4 naïve observers who did not participate in the experiment (600 trials each, 25 face identities). We selected 15 face identities based on mean response deviating no more than ±10% from target accuracies (75% male response for male faces, 50% male response for androgynous, 25% male response for female). The phase-resetting auditory stimulus was a 16 ms, 900 Hz pure tone (80 dB sound pressure level at the ear, 44100 kHz sampling frequency) projected through 2 loudspeakers besides the monitor (following Romei *et al.* [37]).

### Procedure

Participants fixated a white fixation dot at the center of the screen, which was present for the whole duration of the experiment except during face stimulus presentation. Each trial began with the presentation of the auditory stimulus. After an interval ranging uniformly from 100 to 1000 ms (at 120 Hz sampling frequency, the monitor refresh rate) one of the 45 faces was presented for 17 ms (two frames). Participants were instructed to wait at least 1 second before responding, indicating whether the presented face seemed male or female (by pressing the left or right arrow keyboard keys). Trials where participants responded earlier than 1 second were eliminated from the analysis. Response configuration (association of arrows with gender) was randomized between participants, and switched halfway through the experiment. After button press, a new trial started after an interval ranging uniformly from 800 to 1100 ms, so that the auditory stimulus presentation was not easily predictable. Each participant completed 1215 trials.

### EEG Acquisition and Preprocessing

EEG data were collected with a Nautilus Research headset (g.tec) at a sample rate of 250 Hz with no online filtering. The data were referenced online to a unilateral electrode placed behind the left ear. Activity was measured from 32 gel-based active electrodes (g.LADYbird technology) arranged according to the 10/20 system. Impedance was kept below 50 kΩ.

Offline EEG preprocessing was performed in MATLAB (MathWorks®) with custom code. EEG data were referenced to the common average reference and filtered with a FIR bandpass filter (Chebyshev window, 128^th^ order, stopbands 1 Hz and 35 Hz, sidelobe magnitude factor 50 dB). For the main data analysis, epochs were extracted aligned to stimulus presentation, comprising a segment of data from −500 ms to 1800 ms after the stimulus. Epochs were visually inspected for motor artifacts and wireless failure of signal transmission (manual rejection of 0.8% ± 0.6% STD of data across subjects). Ocular artifacts were removed through blind source separation with ICA decomposition [58].

### Data analysis *–* psychophysics

We analyzed individual responses to androgynous face identities, calculating the proportion of “male” responses, depending on the response to the previous trial. We removed from all analyses trials where response latencies were lower than 1 second or higher than 3 seconds (average response latency 2.2 s; CI95 = 1.6–2.7 s). After artifact removal we obtained 1249 ± 9 trials per subject. To assess whether previous responses changed gender discrimination bias, we compared by t-test proportion “male” responses when previous response was male to when previous response was female (only on androgynous trials). The same analysis was repeated based on face identity of the previous trial (stimulus-based analysis).

To measure oscillations in face-gender behavioral bias, we applied single-trial analysis to aggregate data from all participants, including all the trials that survived the EEG artefact rejection procedure. The general linear model (GLM) analysis weighted each single trial with the following model:

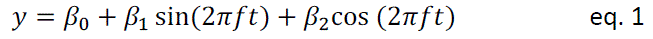

where *y* is participant response (1 for male, 0 for female), *t* is SOA (audio to face) in seconds, *f* is a fixed frequency ranging from 4 to 20 Hz at 0.5 Hz intervals, β0, β1 and β2 are free parameters. We separated responses to all stimuli based on previous response (male or female), and fit the above GLM, calculating the amplitude of the sinusoidal fit to each frequency in the range 10-20 Hz. To assess statistical significance, the surrogate data generated by shuffling the responses (2000 permutations) were analyzed with the same GLM obtaining a distribution of amplitudes across frequencies under the null hypothesis. Frequencies with amplitudes over the 95^th^ percentile of this distribution were candidates for statistical significance. Candidate frequencies were deemed statistically significant only if they were included in a cluster larger than the 95^th^ percentile of the distribution of cluster sizes. For each cluster in both previous male and previous female conditions, we noted the local maxima of amplitudes and their corresponding frequencies. For illustration purpose only (Fig. 6) we show the responses binned on stimulus onset asynchrony (SOA) at 8.3 ms intervals with a running average of 3 consecutive bins (weights: 0.305, 0.39, 0.305).

**Figure 6.**
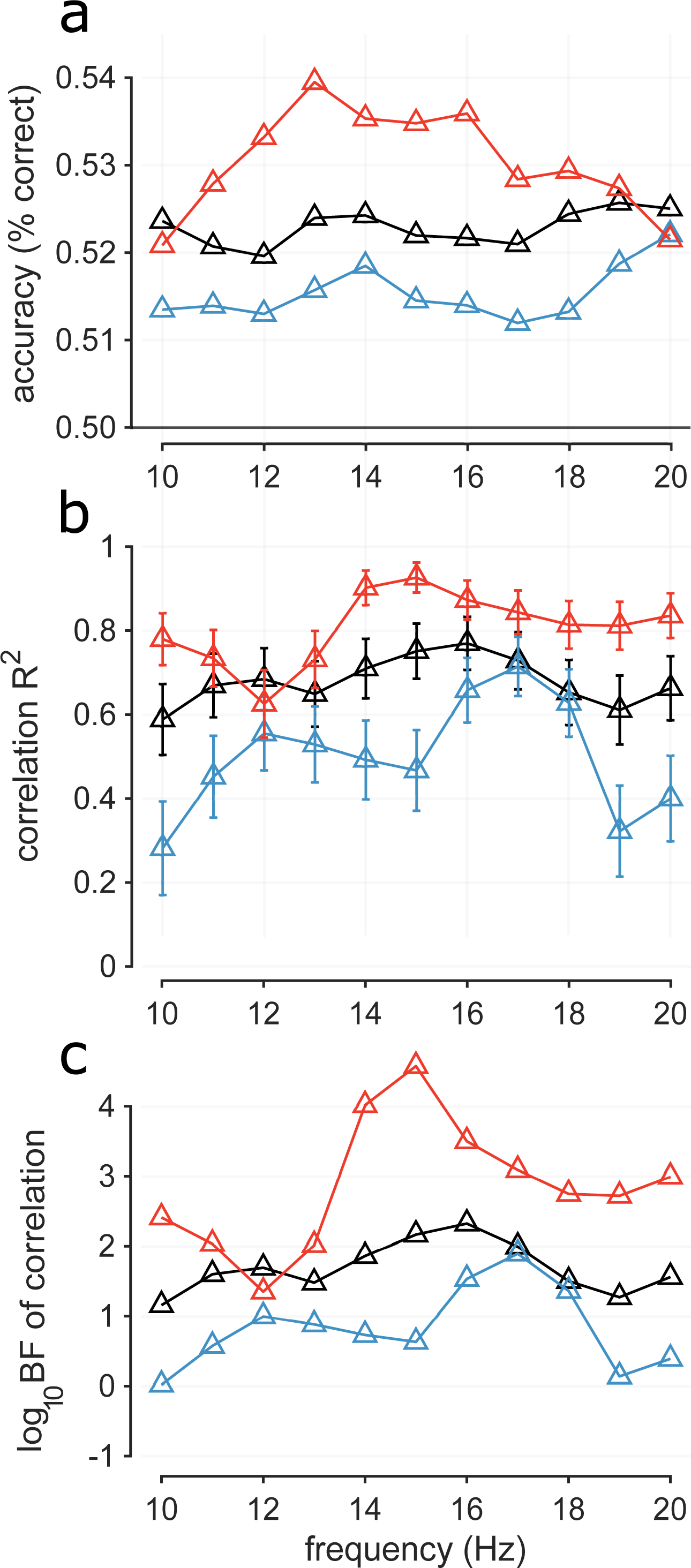
Decoding across narrow frequency ranges. **A.** Classification accuracy across frequency calculated as average across the diagonal ± 2 points (precision ±60 ms) in the window 0–1 s. Classification accuracy tested on all trials (black), on androgynous trials with female response (red) and on androgynous trials with male response (blue). **B.** Squared correlation coefficients (variance explained) between classification accuracy and serial dependence strength across participants. Classification accuracy calculated as in A. Serial dependence effect (constant across all 3 conditions) was calculated as the difference between proportion male response with previous male response and proportion male response with previous female response. **C.** Log10 of bayes factor of the correlations of Fig. B.

### Data analysis – EEG time-domain and power analysis

We calculated event-related potentials (ERPs) synchronized to face stimulus presentation, separately for the two conditions “previous male” and “previous female”. Grand-average ERPs were calculated by averaging single-participant ERPs normalized (z-scores) to a baseline window in the interval from –500 to –100 ms with respect to stimulus presentation.

We classified previous (*n*-1) responses from the EEG activity elicited by the current (*n*) trial. Before classification, we filtered the entire time series of 111 ± 18 minutes across participants, in a low-beta range (14 to 20 Hz) and calculated the amplitude of the analytic envelope using the Hilbert transform (Fig. 3). For the second decoding procedure (Fig. 5) we filtered narrowly around frequencies from 4 to 20 Hz at 1 Hz steps (Type II Chebyshev window, 1024^th^ order, passband 0.5 Hz above and below the selected frequency, stopband 1 Hz above and below the selected frequency, sidelobe magnitude factor 50 dB). We down-sampled trials to 50 Hz, obtaining 150 time points from –500 to 1000 ms, synchronized to face stimulus presentation. To boost signal-to-noise ratio, we averaged 5 trials of the same class together (as in Foster et al., 2016), obtaining our final classification samples. We used binary support vector machine (SVM) classifiers with linear kernel (the MATLAB *fitcsvm* function), using the 32 electrode locations as features for classification. Samples were split in 5 folds, with a 4:1 training to testing ratio (196 ± 14 SD samples in the training set per observer). The procedure was repeated 5 times, rotating which fold was used for testing. The whole decoding procedure was repeated 100 times, generating new samples every time by averaging random sets of 5 trials together. This allowed us to minimize lucky splits of the data and assess accuracy by averaging over a large number of guesses (24,519 ± 141 SD guesses per observer).

We assessed temporal generalization by testing classifiers across all time points, obtaining a matrix of accuracy across training and testing times per participant. We averaged accuracy across participants and extracted matrix diagonals to show the dynamics of previous response signals. Statistical power was defined by t-tests against 50% accuracy across training and testing time. To correct for multiple comparisons, we permuted class labels at the level of testing, obtaining a set of 2000 temporal generalization matrices under the null hypothesis. For each matrix, we noted how many adjacent points survived a t-test against 50% accuracy, generating a distribution of cluster sizes. Calculating the 95^th^ percentile of this distribution, we obtained a threshold of cluster size. All clusters smaller than the identified thresholds were deemed non-significant.

Activation patterns show the relative relevance of electrode sites for classification, giving qualitative insight on model dynamics. Activation patterns were calculated as follows:

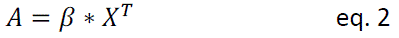

Where *β* are classifier coefficients and *X^T^* is the transposed matrix of EEG signal of tested data.

Classifiers trained on all trials were then tested on 2 subsets of trials: androgynous trials to which observers responded “male” and androgynous trials which observers responded “female”. Class labels within these conditions were again responses to the previous trial. As trials in the two conditions are likely trials where serial biases were more present, this testing procedure highlighted whether the information is stronger in male-biased or female-biased trials.

